# Mitochondriomics reveals the underlying neuoprotective mechanism of TrkB receptor agonist R13 in the 5×FAD mice

**DOI:** 10.1101/2021.02.08.430227

**Authors:** Xiao Li, Ting Li, Hao Yu, Shupeng Li, Zaijun Zhang, Yongmei Xie, Xiangrong Song, Jianjun Liu, Xifei Yang, Gongping Liu

**Author notes:** Address correspondence to: Dr. Xifei Yang, Key Laboratory of Modern Toxicology of Shenzhen, Shenzhen Center for Disease Control and Prevention, No. 8, Longyuan Road, Nanshan District, Shenzhen, China 518055.; Dr. Gong-Ping Liu Department of Pathophysiology, School of Basic Medicine and the Collaborative Innovation Center for Brain Science, Key Laboratory of Ministry of Education of China and Hubei Province for Neurological Disorders, Tongji Medical College, Huazhong University of Science and Technology, Wuhan 430030, China. These authors contributed equally to this work (Xiao Li, Ting Li).

## Abstract

Decreased energy metabolism and mitochondrial biogenesis defects are implicated in the pathogenesis of Alzheimer’s disease (AD). In present study, mitochondriomics analysis revealed significant effects of R13, a prodrug of 7,8-dihydroxyflavone, on mitochondrial protein expression profile, including the proteins related to the biological processes: fatty acid beta-oxidation, fatty acid metabolic process, mitochondrial electron transport, and mitochondrial respiratory chain. Cluster analysis of mitochondriomics demonstrated that R13 promoted mitochondrial oxidative phosphorylation (OXPHOS). The functional analysis showed that R13 increased ATP levels and enhanced OXPHOS including complex I, complex II, complex III and complex IV. R13 treatment increased mitochondrial biogenesis by regulating the levels of p-AMPKα, p-CREB, PGC-1α, NRF1 and TFAM as a consequence of activation of TrkB receptor in the 5×FAD mice. Finally, R13 significantly reduced the levels of tau phosphorylation and Aβ plaque. Our data suggest that R13 may be used for treating AD via enhancing mitochondrial biogenesis and metabolism.

## Introduction

Alzheimer’s disease (AD) is the most common type of dementia, mainly characterized by Aβ amyloid deposits and neurofibrillary tangles in the brain. It was reported that the number of dementia patients worldwide reached 24.3 million in 2001 and may be further expanded to 42.3 million in 2020 and twice as many as 2020 by 2040 (Ballard, Gauthier et al. 2011). AD is divided into two types: sporadic AD and familial AD (FAD). Approximate 50% of FAD patients show autosomal dominant inheritance of mutations including amyloid precursor protein (APP), presenilin-2 (PS2), and presenilin-1 (PS1) (Villamil-Ortiz, Barrera-Ocampo et al. 2018). However, the specific molecular mechanism of AD is still unclear. In order to explore the pathological process of AD, various animal models have been established for experimental research, among which 5×FAD mice is one of the most widely used. 5×FAD mice contain human amyloid beta precursor protein (APP) with three FAD mutations (K670N/M671L, V717I and I716V) and Presenilin (PS1) with two FAD mutations (L286V and M146L). Amyloid deposition of 5×FAD begins at 2 months and increases with age in mice brain. Synapse-associated protein including synaptophysin, syntaxin and postsynaptic density-95 (PSD95) reduce with age in the brain of 5×FAD mice (Oakley, Cole et al. 2006).

Mitochondria are important organelles that provide energy for brain. The increasing evidence shows that mitochondrial dysfunction is related to many neurodegenerative diseases, including AD, amyotrophic lateral sclerosis, Parkinson’s disease and Huntington’s disease (Bender, Krishnan et al. 2006, Zhu, Perry et al. 2006, Reddy, Mao et al. 2009, Ferraiuolo, Kirby et al. 2011). In fact, mitochondrial damage is important and early features of AD with declined energy metabolism (Sheng, Wang et al. 2012). Mitochondrial biogenesis is disturbed in an AD model, in which PPARγ-coactivator-1α (PGC-1α) was significantly reduced (Katsouri, Lim et al. 2016). Therefore, mitochondrial dysfunction leads to reduced ATP levels, which would aggravate mitochondrial damage, and promotes the progression of AD (Mattson 2004). In addition, mitochondrial DNA (mtDNA) defects and mutation are also found to be associated with the incidence of AD (Lin, Simon et al. 2002).

Currently, drugs that be clinically used to treat AD such as memantine, donepezil, galantamine, and rivastigmine, only relieve the symptoms of AD, but are unable to cure or delay AD. Therefore, it is imminent to discover new drugs for AD. Brain-derived neurotrophic factor (BDNF) is an important neurotrophic factor, which regulates neuronal development, differentiation and survival by activating TrkB receptor (Liu, Wei et al. 2020). The expression level of BDNF is decreased in the brain of AD patients (Zuccato and Cattaneo 2009). Moreover, BDNF has been used as a therapeutic target in various AD mouse models (Nagahara, Merrill et al. 2009). BDNF exerts physiological functions by TrkB receptor and its downstream signaling pathways involving MAPK-ERK (extracellular signal-regulated kinase), the phospholipase Cγ1 (PLCγ1)-PKC pathway and PI3K-AKT pathway (Longo and Massa 2013). However, the failure of clinical trials of recombinant BDNF may be due to the short in vivo half-life of BDNF (Chen, Wang et al. 2018). Therefore, the idea to find a chemical compound to mimic the effect of BDNF is still an attractive option for AD therapy.

In this work, we found that R13, a prodrug of 7,8-DHF that acts as a TrkB receptor agonist, improved learning and memory in 5×FAD mice by activating TrkB receptor. By using proteomics, we drew the heatmap of 139 differentially expressed mitochondrial proteins in the hippocampus and found that R13 treatment significantly enhanced oxidative phosphorylation (OXPHOS) of 5×FAD mice. R13 enhanced the ATP levels in the hippocampus of 5×FAD mice by increasing proteins levels involving in OXPHOS. Moreover, R13 promoted mitochondrial biogenesis and increased mitochondrial copy number. Finally, the levels of tau phosphorylation and Aβ plaque were significantly decreased in R13-treated 5×FAD model.

## Materials and methods

### Animals and treatments

The 5×FAD mice (expressing APP_K670N/M671L_, APP_V717I_, APP_I716V_, PS1_L286V_ and PS1_M146L_) were obtained from the Jackson Laboratory (Maine, USA). The 5×FAD mice were housed in per cages (5 mice per cage) and maintained in a 12-h light-dark cycle-controlled environment with free food and water. All animal experiments were performed following the ’Policies on the Use of Animals and Humans in Neuroscience Research’ revised and approved by the Society for Neuroscience in 1995. This study was approved by the Ethics Committee of the Shenzhen Center for Disease Control and Prevention.

R13 compound was dissolved in 5% DMSO/0.5% methylcellulose (pH=2) (Chen, Wang et al. 2018). The 2-m-old 5×FAD mice were orally treated with vehicle or R13 for 3 consecutive months at a dosage of 3.6, 21.8 or 43.6 mg/kg/d, respectively. After the completion of behavioral tests, brain tissues were removed and the bilateral hippocampus was isolated and stored at -80 °C for future analysis.

### Proteomics

#### Preparation of mitochondria of hippocampal tissue samples

Mitochondria of hippocampal tissue were isolated using the Qproteome Mitochondria Isolation Kit (Qiagen 37612, Germany). Mitochondria of hippocampal tissue (3 mice each group) placed in 8M urea dissolved in PBS were lysed with ultrasonic disruption for protein extraction. Then, sample lysates were placed on ice for 30 min and centrifuged at 12, 000g, for 20 min at 4 °C, and the supernatant was used for proteomics. The protein concentration was measured with the BCA protein assay kit (Thermo Fisher, NJ, USA).

### TMT labeling

The TMT labeling of peptide was based on the method reported in the previous published paper (Chen, Jiang et al. 2019). Specifically, the sample with 50 µg protein was treated with 10 mM dithiothreitol at 55 °C for 1 h, and t hen incubated with 25 mM iodoacetamide for 1 h at room temperature in a dark place. Then, these samples were diluted with PBS (pH=8.0) (final concentration: 1 M urea), digested with trypsin (1:25w/w) (Promega, WI, USA) for 15 h at 37 °C. After digestion, peptides were adjusted to pH=2 with 0.1% trifluoroacetic acid to stop reaction and then centrifuged at 12,000 g for 10 min to collect the supernatant. The supernatants were desalted with reversed-phase column (Oasis HLB; Waters, Milford, MA, USA), dried with vacuum concentrator, and then dissolved in 50 μl triethylammonium bicarbonate (TEAB, 200 mM). Peptides were labeled with the TMT reagents at room temperature for 1 h: TMT-126 for mitochondrial proteins of the Wild type (Wt), TMT-127 for mitochondrial proteins of the 5×FAD, TMT-128 for mitochondrial protein of the 3.6 mg/kg R13-treated 5×FAD, TMT-129 for mitochondrial proteins of the 21.8 mg/kg R13-treated 5×FAD, TMT-130 for mitochondrial proteins of the 43.6 mg/kg R13-treated 5×FAD . The labeling reaction is terminated by 5% hydroxylamine. The labeled peptides were mixed (n=3), desalted, dried, and dissolved in 100 µL 0.1% FA.

### High performance liquid chromatography (HPLC) separation

The labeled peptides were fractionated according to methods in our lab (Gokce, Andrews et al. 2011). Briefly, these peptides were loaded onto the Xbridge BEH300 C_18_ column (Waters Milford, MA, USA) for HPLC (UltiMate 3000 UHPLC; Thermo Fisher Scientific, Waltham, MA, USA) separation. The fractions were collected from 3 min to 75 min during separation, then dried, finally dissolved in 20 µl 0.1% FA for further liquid chromatography (LC)-mass spectrometry (MS)/MS analysis.

### Peptide analysis by LC-MS/MS and database searching

The peptide analysis was conducted by LC-MS/MS using a previously described procedure (Xu, Zhang et al. 2018). Brief, the peptides were separated by liquid chromatography and analyzed by Q-Exactive mass spectrometer (Thermo Scientific, NJ). Original mass spectrum data was searched against the UniProt-Mus musculus database (released on November 07, 2019) using Proteome Discoverer 2.1 software (Thermo Scientific). Differential expression (DE) proteins are defined according to a standard (P < 0.05).

### Bioinformatics analysis

We firstly used Perseus software to screen out the DE proteins in 5×FAD model and the R13-treated 5×FAD . Then, by using cellular components of GO terms and uniport databases, 139 DE proteins were localized in mitochondria. We have completed the biological process analysis in Database for Annotation, Visualization and Integrated Discovery (DAVID). We used GraphPad Prism 8.0 to draw the mitochondrial heatmap of these proteins and biological process. Cluster analysis of these proteins was performed by RStudio. We performed Molecular Complex Detection (MCODE) and Clue GO analysis by using cytoscape software (3.7.1). The pathways of three clusters were carried out in the website (http://metascape.org/gp/index.html#/main/step1).

### Western blotting

The hippocampus was lysed ultrasonically in the RIPA (Beyotime, P0013B) containing protease and phosphatase inhibitor (Thermo Scientific, NJ, USA). Hippocampal protein concentration was measured by NanoDrop 2000/2000c spectrophotometers (Thermo Fisher Scientific, Waltham, MA, USA). The proteins were separated by 8%∼12% SDS-PAGE and transferred to polyvinylidene fluoride membranes. Membranes were blocked with 5% skimmed milk in TBST buffer for two hours at room temperature, and then incubated with the following primary antibodies in the TBST buffer at 4 °C overnight. The application information of primary antibodies was as follows: COX5B (Abcam, ab180136), NDUFA10 (Abcam, ab103026), SDHB (Abcam, ab14714), UQCRFS1 (Abcam, ab131152), ATP5A (Abcam, ab14748), TFAM (Abcam, ab131607), NRF1 (Abcam, ab175932), PGC-1α (Abcam, ab54481), Cytochrome c (Cell Signalling, #11940), VDAC1 (Abcam, ab154856), PDH (Cell Signalling, #3205), AMPKα (Cell Signalling, #2532), p-AMPKα (Cell Signalling, #2535), CREB (Cell Signalling, #9197), p-CREB (Cell Signalling, #9198), Akt (Cell Signalling, #4691), p-Akt (Cell Signalling, #4060), p-ERK (Cell Signalling, #4370), ERK (Proteintech, 16443-1), p-TrkB (Sigma, ABN1381), TrkB (Biovision, #3593), pS396 (Abcam, ab109390), pS404 (Abcam, ab926736), Tau 5 (Abcam, ab80579), Tubulin (Merck, MAB1637), β-actin (Santa cruz, 47778). After washing three times in TBST for 10 min each, the membranes were incubated with a secondary antibody (1:10,000) at room temperature for 1 h. Then, the membranes were washed three times in TBST for 10 min each. Finally, the membranes were exposured by the ECL kit.

For dot blot, we added the same amount of cortical protein to the nitrocellulose membrane, and let the membrane dry. Block non-specific sites by soaking in 5% BSA in TBST in a 10 cm Petri dish. The following steps are similar to western blotting.

### Hippocampal ATP level assessment

We used an ATP Assay Kit (Beyotime, S0026) to measure the ATP levels in hippocampal tissues. Shortly, hippocampal tissue was homogenized in ATP lysate, and then centrifuged to preserve the supernatant. Protein concentration of the supernatant was determined by the BCA method. Hippocampal sample (20 μl) and ATP detection working solution (100 μl) were added to the detection hole, and incubated for 3 min. ATP level was detected with multifunctional microplate reader.

### Relative mtDNA copy number

Relative mtDNA copy number was detected according to a previously described procedure (Foote, Reinhold et al. 2018). The total DNA from hippocampus was extracted with kit (Tian gen, DP304-03). The relative mtDNA copy number level was normalized by the nuclear DNA. QPCR primers employed in the present study are as following: the primers for nuclear control: (F) TTGAGACTGTGATTGGCAATGCCT and (R) CCAGAAATGCTGGGCGTACT; the primers for mtDNA: (F) GCCAGCCTGACCCATAGCCATAAT and (R) GCCGGCTGCGTATTCTACGTTA.

### Morris Water maze

Morris Water Maze (MWM) is a classic method for assessing spatial and learning memory in mice (Feng, Luo et al. 2020). This maze was mainly a circular pool with a diameter of 1.5 m, which can be divided into four quadrants. A platform was placed in one of the quadrants, and then water was poured until the platform was approximately 2 cm submerged and added milk powder to the water. In this study, the platform was put in the third quadrant. The training period was 5-consecutive days, and the main task was to train mice to find the hidden platform. Mice were placed into the water with facing the wall, and placed once in each quadrant in a random order, four times a day, with an interval of 1 h. If mice find the platform within 60s, the time it took to find the platform was the escape latency. If the platform was not found in 60s, the escape latency was recorded as 60s and mice were guided onto the platform and stayed for another 30s. Memory test was performed at 1 day after the training. During the test, the platform was removed, and the following contents were mainly recorded: the escape latency, the time spent in the quadrant where the platform was located previously, and the number of times mice crossed the place where the platform used to be. All the behaviors of mice in the experiment were monitored by the camera above the pool, which connected to a digital-tracking device linked to a computer, and all data were analyzed by software.

### Immunohistochemistry

Abcam ABC HRP Kit (ab64264) was used and all procedures were carried out according to the instructions. Specifically, paraffin-embedded brain slices were firstly deparaffinized in xylene and then rehydrated in gradient alcohol. The brain slices in 10 mM citric acid are heated in a microwave oven on medium heat for 10 min, and then placed at room temperature to cool for 30 min for antigen retrieval. Then, enough drops of 3% H_2_O_2_ were added to cover the sections for 10 min followed by washing 3 times in PBS for 5 min each time. Then, 3% BSA was dropped on brain slice to incubate for 1 h at room temperature. Samples were incubated with Anti-β-Amyloid (#803015, Biolegend) at 4 °C overnight, followed by incubation for 10 min with biotinylated goat anti-polyvalentand and streptavidin peroxidase at room temperature. Then, brain slices were stained for 2 min using diaminobenzidine (DAB). After dehydrating with gradient alcohol, the sections were immersed in xylene for transparency, and sealed by using neutral resin. Finally, brain slices were imaged with an optical microscope.

### Statistical analysis

All the data were presented as mean ± standard error (S .E.M), and all statistical tests were carried out using two-way repeated measures ANOVA tests or one-way ANOVA followed by Tukey’s multiple comparison tests with the GraphPad Prism 8 (GraphPad Software, USA). P < 0.05 was considered as statistically significant difference.

## Results

### R13 improved cognitive deficits of 5×FAD mice

As previous described, 5×FAD mice showed obvious impairments of learning and memory at 4∼5 months old (Oakley, Cole et al. 2006). We performed MWM test to explore the effect of R13 on cognitive functions of 5×FAD model after 3 months of oral administration. In training stage, the 5×FAD mice spent more time to find the submerged platform compared with the wild-type littermates (at 2^nd^ and 5^th^ d) (Fig. 1A), while R13 treatment obviously decreased the escape time of 5×FAD mice in the dose of 21.8 mg/kg (at day 5^th^ d) or 43.6 mg/kg (3^rd^, 4^th^ and 5^th^ d) (Fig. 1A), which suggested that R13 ameliorated the learning abilities. Moreover, chronic oral 21.8 or 43.6 mg/kg R13 decreased the latency to find the submerged platform during the test stage, which indicated that R13 restored the memory of 5×FAD mice (Fig. 1B). According to the molecular weight of R13 and 7,8-DHF, 15 mg/kg of 7,8-DHF are equivalent to 21.8 mg/kg of R13. The improved performance of 5×FAD during training implied that R13 has better therapeutic effect than corresponding dose of 7,8-DHF. In the previous study, administration of R13 releases a higher concentration of 7,8-DHF in the brain than direct administration of 7,8-DHF (Chen, Wang et al. 2018). There was no significant difference in the number of platform crossing, time spent on the target quadrant, swimming speed or distance among 6 groups (Fig. 1C-F). These data demonstrate that R13 reverses the spatial learning and memory impairments of 5×FAD mice.

**Fig 1.**
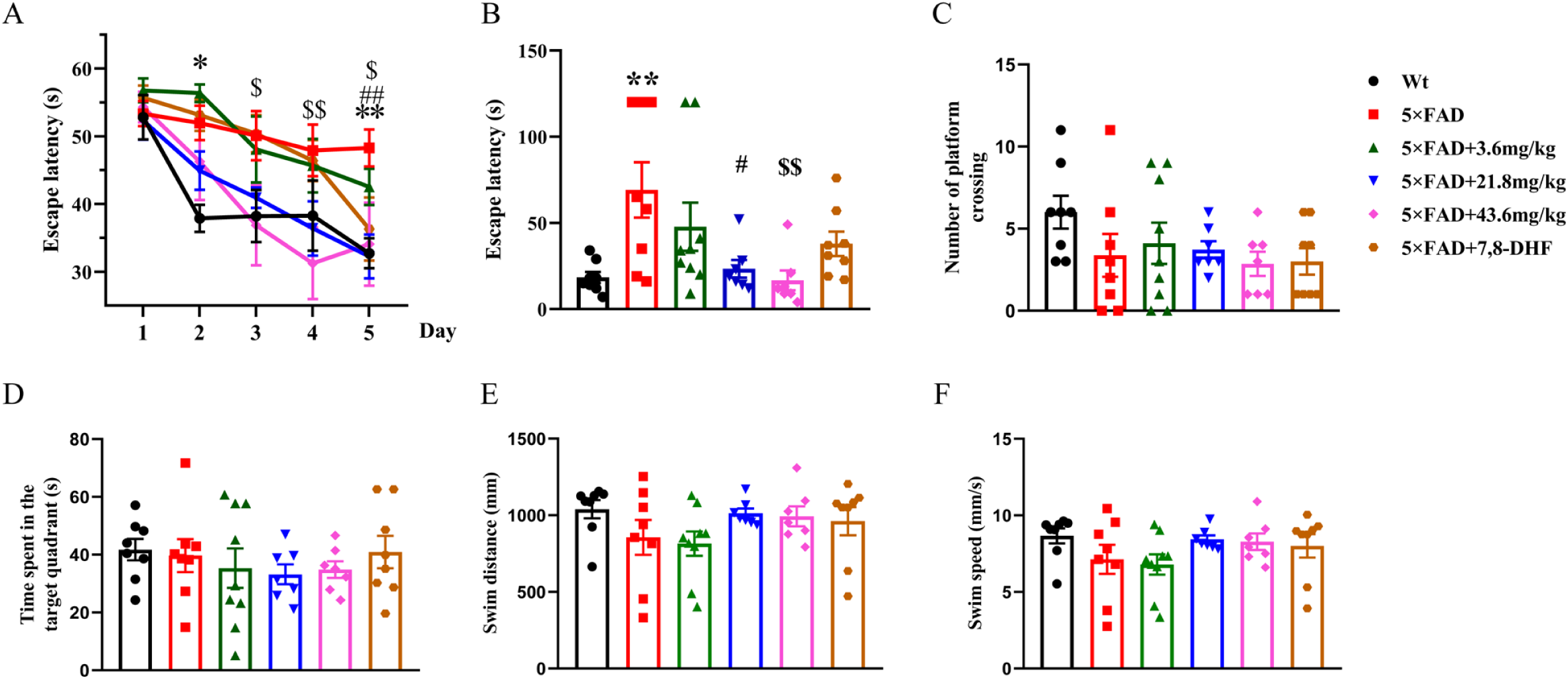
R13 treatment reversed learning and memory defects of 5×FAD mice. (A) During training test, R13 treatment improved 5×FAD mice’s learning ability, and (B) the memory capacity, which was shown by the obviously decreased escape latency detected by MWM. (C, D) There was no significant difference in number of platform crossing and time spent in the target quadrant during test among 6 groups. (E, F) There was no significant difference in swimming speed or swimming distance among the mice of 6 groups. All data were expressed as mean ± S.E.M., N=7-9 mice for each group. *, *p* < 0.05, **, *p* < 0.01 (5×FAD +vehicle *vs* WT); ##, *p* < 0.01 (5×FAD +21.8mg/kg *vs* 5×FAD +vehicle); $, *p* < 0.05, $$, *p* < 0.01 (5×FAD +43.6mg/kg *vs* 5×FAD +vehicle).

### Mitochondriomics profile in R13-treated 5×FAD mice

In the present study, we used the six TMT method and two-dimensional LC/LC-MS/MS (Guo, Pang et al. 2019), and then conducted the mitochondrial proteome in five groups excluding the 7,8-DHF-treated group (Fig. S1). We detected 4999 proteins in the proteome (false discovery rate (FDR)< 1%), of which there were 139 differentially expressed (DE) proteins (p value < 0.05) localized in the mitochondria.

To investigate the effect of R13 on the mitochondria of 5×FAD mice, we divided these DE proteins localized in the mitochondria into 10 types by using GO terms and uniport databases, and accomplished mitochondrial protein heatmap (Fig. 2A). Groups 0, 1, 2, 3, 4 respectively correspond to Wt, 5×FAD, 5×FAD +3.6 mg/kg, 5×FAD +21.8 mg/kg, 5×FAD +43.6 mg/kg. These 10 types included: mitochondrial out membrane, mitochondrial inner membrane, mitochondrial intermembrane space, mitochondrial matrix, oxidative phosphorylation (OXPHOS), ribosomal protein, apoptotic process, ATP binding, metabolic process and others.

**Fig 2.**
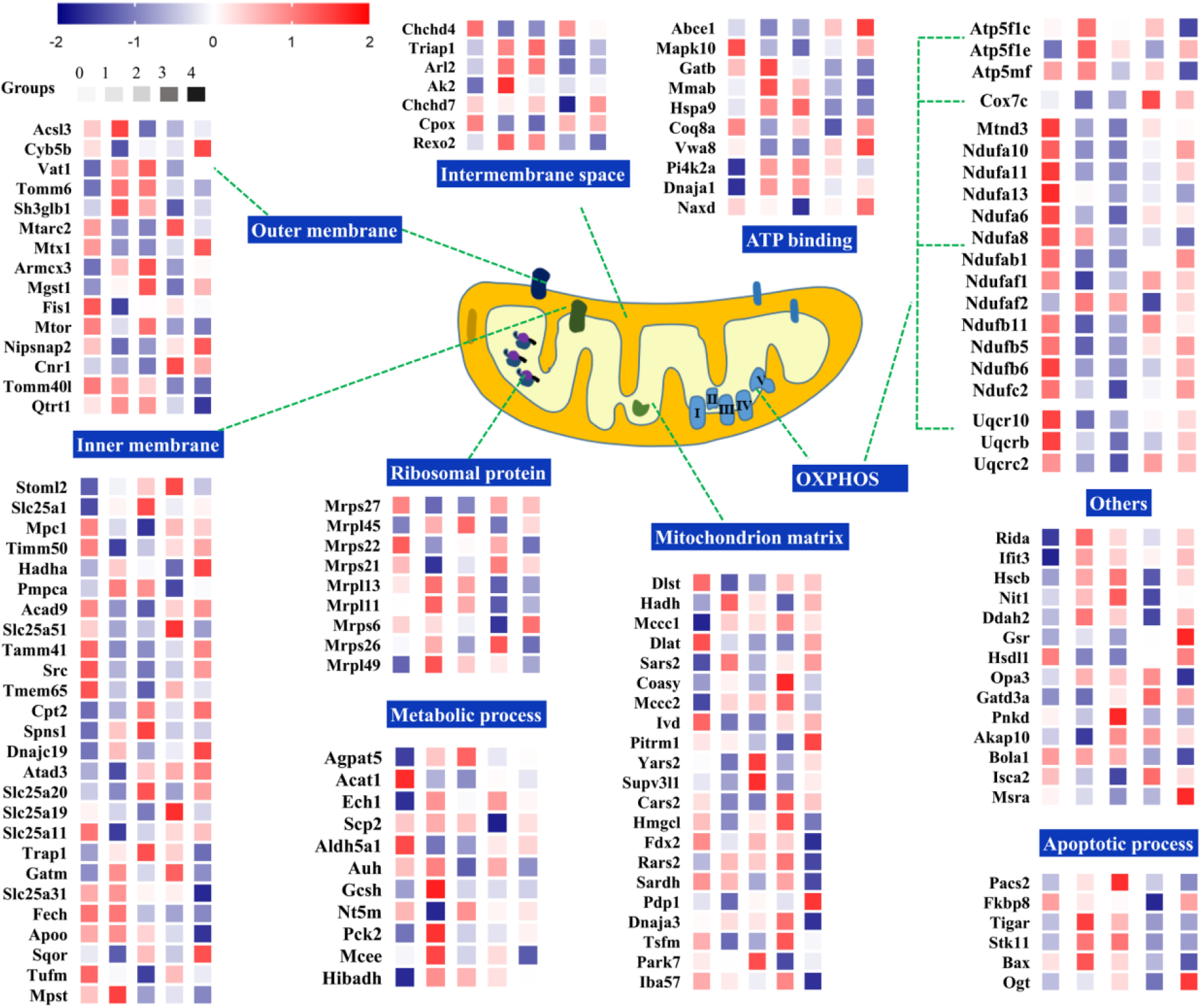
Heatmap of DE protein localized in mitochondria. Groups 0-4 correspond to the group of Wt, 5×FAD, 5×FAD + 3.6 mg/kg, 5×FAD + 21.8 mg/kg, or 5×FAD + 43.6 mg/kg, respectively. The average of each protein in the three independent mice is standardized on the wukong website (https://www.omicsolution.org/wkomics/main/). And then these data were used to draw the mitochondrial heatmap. Red and blue indicate up or down regulation, respectively (N=3 mice/per group).

As shown in the heatmap, complex I was the most widely affected by R13 treatment relative to other complexes. The subunit proteins of complex I were mostly decreased except for ndufaf2 in 5×FAD model, and significantly upregulated in oral R13-treated 5×FAD model mice. Moreover, the change of complex III containing uqcr10, uqcrb, and uqcrc2 was similar to complex I. Complex V involved Atp5flc, Atp5fle, Atp5mf, was increased in model compared to Wt, while their levels were restored by R13 treatment. The level of Cox7c (Complex IV) was decreased in model, which was significantly upregulated at 21.8 mg/kg R13-treated 5×FAD mice. These results indicate that complex I, complex III and complex IV are markedly improved by oral R13 administration. Although Complex V was downregulated by R13 treatment, R13 treatment restored the level of complex V to that of Wt mice.

Then, we performed cluster analysis of these DE proteins, which were divided into three categories (Fig. 3A). By MCODE analysis, Cluster 1, 2, or 3 was related to translation, OXPHOS, or fatty acid oxidation (FAO), respectively (Fig. 3B). Cluster 1 (n=57) displayed an apparent increase in 5×FAD mice compared with Wt mice, but these proteins were slightly decreased by R13 treatment (Fig. 3A). Cluster 1 contained some biological process: negative regulation of protein ubiquitination, fatty acid beta-oxidation, fatty acid metabolic process, metabolic process, leucine catabolic process, oxidation-reduction process, protein oligomerization, and response to insulin (Fig. 3C). These proteins in the cluster 1 were enriched in 8 pathways (Fig. 3D). Similarly, proteins in the cluster 2 (n=47) had the opposite changed trend compared to cluster 1. Furthermore, the 21.8 mg/kg dose displayed the best reversed-effect on protein expression compared to other doses (Fig. 3A). These proteins included biological processes as follows: oxidation-reduction process, transport, mitochondrial electron transport, mitochondrial respiratory chain complex I assembly, translation, response to oxidative stress, aerobic respiration, metabolic process, mitochondrial acetyl-CoA biosynthetic process from pyruvate, and mitochondrial respiratory chain complex III assembly (Fig. 3C). Cluster 2 contained four pathways, which was the mainly relevant to TCA and respiratory electron transport (Fig. 3D). Finally, the changing trends of proteins levels in the cluster 3 (n=35) were similar to cluster 1, and there was a significant dose-dependent effect at 21.8 mg/kg and 43.6 mg/kg (Fig. 3A). These proteins involved some biological processes as follows: translation, mitochondrion morphogenesis, response to nutrient, negative regulation of endoplasmic reticulum calcium ion concentration, very-low-density lipoprotein particle assembly, mitochondrial fragmentation involved in apoptotic process, and protein folding (Fig. 3C). These proteins of the cluster 3 were enriched in 3 pathways (Fig. 3D). There are 37 overlap proteins among the changed proteins of 5×FAD *vs* Wt and 5×FAD *vs* R13-treated 5×FAD (3.6, 21.8 and 43.6 mg/kg) (Fig. 3E). They were mainly related to the OXPHOS signaling pathway (Fig. 3F). It was previously found that FAO, OXPHOS are key pathways involved in energy metabolism (Wang, Mohsen et al. 2010, Nsiah-Sefaa and McKenzie 2016). Proteomics results indicated that R13 accelerated OXPHOS in 5×FAD model, and improved energy metabolism in 5×FAD mice.

**Fig 3.**
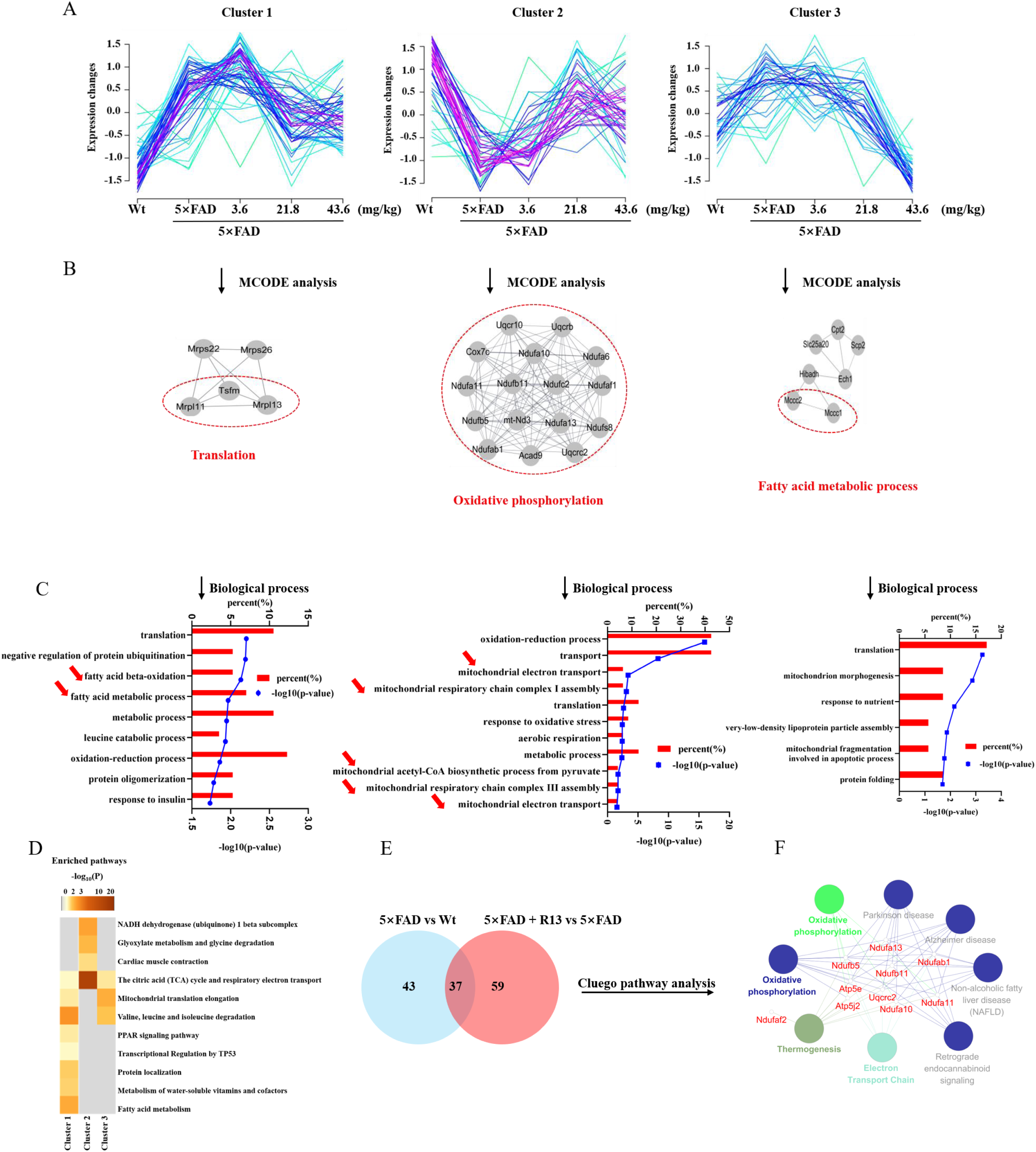
Mitochondriomics analysis in R13-treated 5×FAD mice. (A) DE proteins localized in mitochondria were clustered into three categories by RStudio software. (B) Three types of proteins are analyzed by MCODE in cytoscape software. (C) Enrichment analysis by biological process of 3 clusters was accomplished in DAVID functional enrichment analysis. Red arrows indicated biological processes related to energy metabolism. (D) Enriched pathways in each cluster by metascape website. Significantly enriched pathways (P value < 0.05) are shown. (E) Venn diagram of 5×FAD model and R13-treated 5×FAD changed proteins. (F) Overlap proteins in the Venn diagram (E) were analyzed by ClueGO in Cytoscape software. Significantly enriched pathways (P value < 0.05) are shown.

### R13 restored ATP levels in the hippocampus of 5×FAD mice by enhancing oxidative phosphorylation

It was reported that, accumulation of Aβ damaged mitochondria ATP production (Xu, Mei et al. 2018). Therefore, we assessed ATP levels in the hippocampus of 5×FAD mice and found that ATP levels of 5×FAD mice were significantly decreased compared with the wild-type littermates (Fig. 4A). Strikingly, the decreased ATP levels were noticeably reversed by R13 treatment (Fig. 4A). ATP is mainly produced in the mitochondria through OXPHOS. Therefore, we further detected related mitochondrial OXPHOS protein levels in the hippocampus of 5×FAD mice by immunoblotting. The results showed the complex I (NDUFA10), complex II (SDHB) and complex III (UQCRFS1) levels in the hippocampus of 5×FAD mice apparently decreased, while R13 treatment rescued the reduction of NDUFA10, SDHB and UQCRFS1 (Fig. 4B, C). Although complex V (ATP5A) was not markedly decreased in the model, R13 treatment notably enhanced its protein expression (Fig. 4B, C). Additionally, there was no difference in complex IV (COX5B), but it had a tendency to be increased by chronic oral administration of R13 (Fig. 4B, C). These results were consistent with the proteomics data. In general, these findings suggest that 5×FAD mice had insufficient ATP level, and R13 may increase ATP production by enhancing OXPHOS, thereby reversing the energy deficiency of 5×FAD mice.

**Fig 4.**
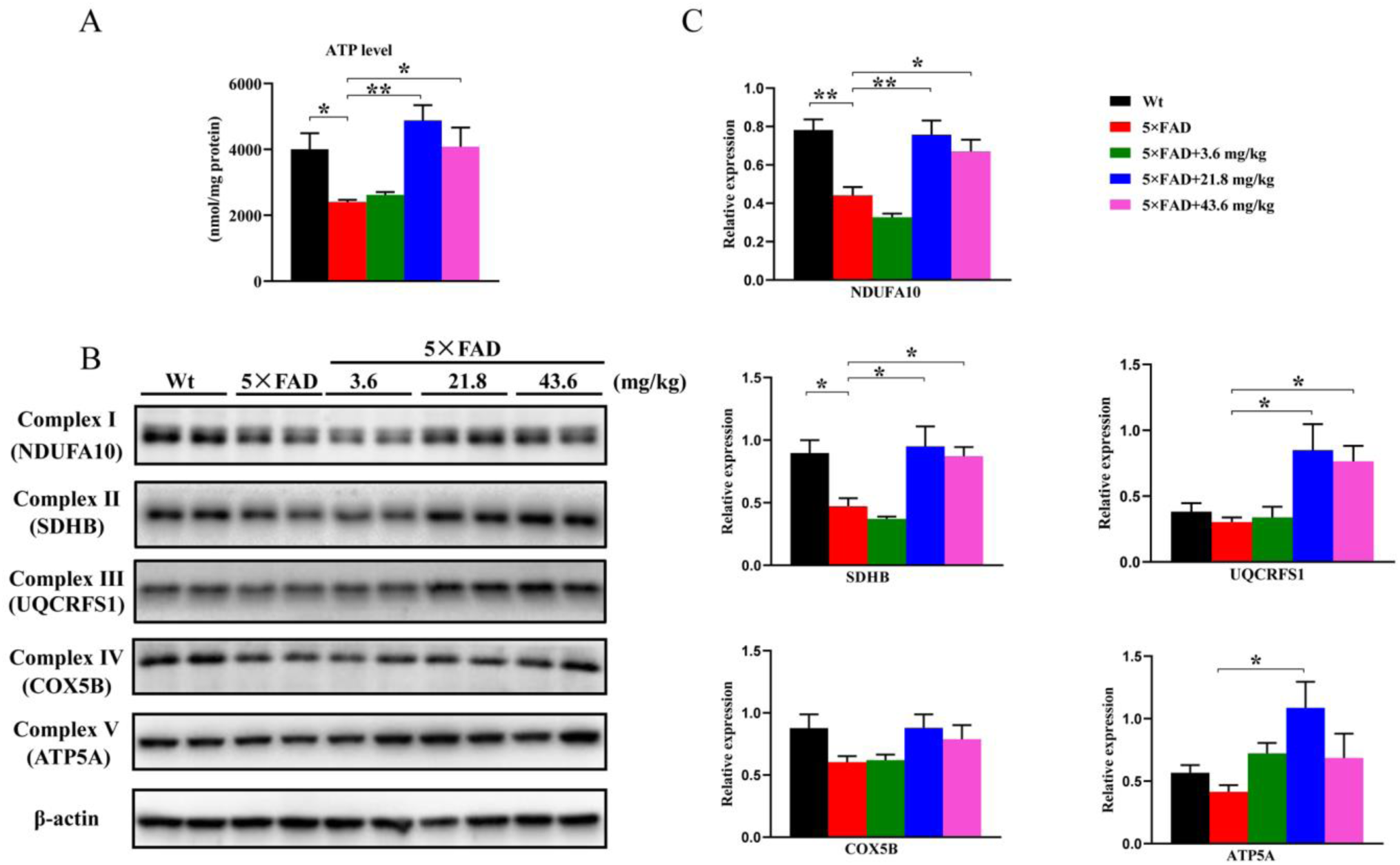
R13 upregulated ATP level by enhancing OXPHOS. (A) The ATP level of 5×FAD mice was decreased compared with Wt mice, which was restored by R13 treatment. (B-C) The relative levels of OXPHOS related proteins were detected by western blotting (B) and quantitative analyses (C). *, *p* < 0.05, **, *p* < 0.01. All data were expressed as mean ± S.E.M (n=4 mice/per group).

### R13 treatment increased mitochondrial biogenesis in the hippocampus of 5×FAD mice

As it is known, mitochondrial biogenesis plays a key role in the formation and maintenance of hippocampal dendritic spines and synapses, and BDNF upregulates PGC-1α levels via CREB (Cheng, Wan et al. 2012). To investigate the molecular mechanisms of R13 improving learning and memory in 5×FAD mice, we analyzed the signaling pathway of mitochondrial biogenesis after R13 treatment. Firstly, we measured the relative copy number of mtDNA. Compared with Wt, the relative mtDNA was significantly decreased in 5×FAD mice, which was reversed by R13 treatment (Fig. 5A). R13 administration also increased the mitochondrial markers including VDAC1, PDH and cytochrome c (Cyto c) (Fig. 5B, C). Furthermore, the phosphorylated level of AMPK at Thr 172 or CREB at Ser 133 in the hippocampus of 5×FAD mice was increased by R13 (Fig. 5D-F). In addition, R13 promoted the expressions of transcriptional factors PGC-1α, NRF1, and TFAM, which were reported to regulate mitochondrial biogenesis (Fig. 5D, G). These data suggest that R13 treatment promotes the mitochondrial biogenesis via AMPK/PGC-1a pathway in the hippocampus of 5×FAD mice.

**Fig 5.**
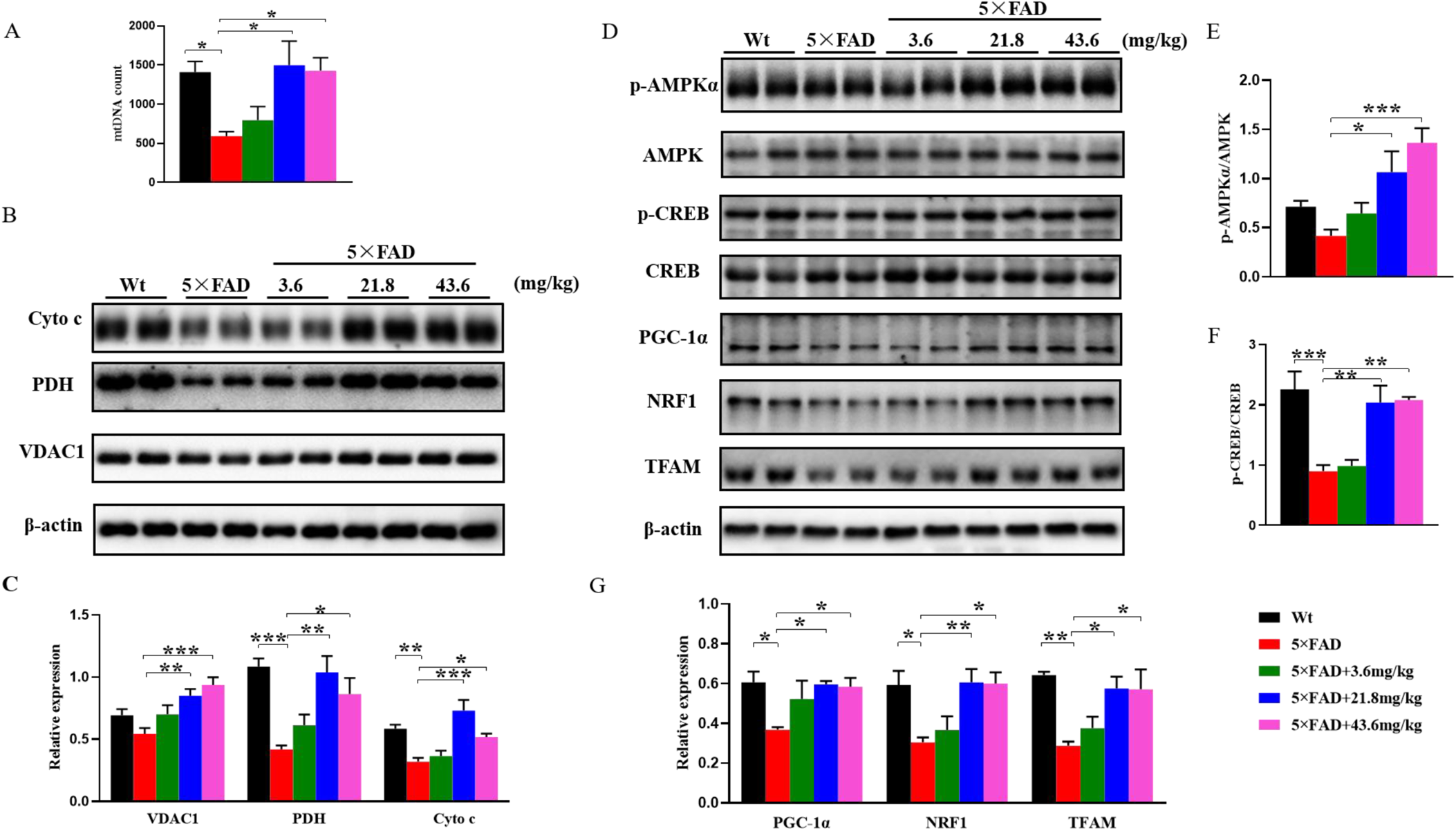
R13 treatment promoted mitochondrial biogenesis. (A) The relative mitochondrial mtDNA copy numbers were quantified by qPCR. (B-C) The mitochondrial markers including Cyto c, PDH and VDAC1 were detected by western blotting (B) and quantitative analyses (C). (D-G) The levels of proteins (p-AMPKα, AMPK, p-CERB, CREB, PGC-1α, NRF1 and TFAM) which regulating mitochondrial biogenesis were detected by western blotting (D), and quantitative analysis (E-G). *, *p* < 0.05, **, *p* < 0.01, ***, *p* < 0.001. All data were expressed as mean ± S.E.M (n=4 mice /per group).

### R13 activates TrkB signaling pathways in 5×FAD mice

To explore whether R13 mimicked the functions of BDNF, we firstly detected the phosphorylated level (p-TrkB, activated form) of TrkB in 5×FAD mice. As expected, the level of p-TrkB relative to total TrkB was markedly increased in R13-treated 5×FAD mice (Fig. 6A-B). Similarly, the downstream signaling molecules AKT and ERK were obviously activated (phosphorylated level increased) in the R13-treated 5×FAD mice than in those treated with vehicle control (Fig. 6A, C, D). Hence, our data suggest that R13 activates TrkB receptor and its downstream signaling pathway.

**Fig 6.**
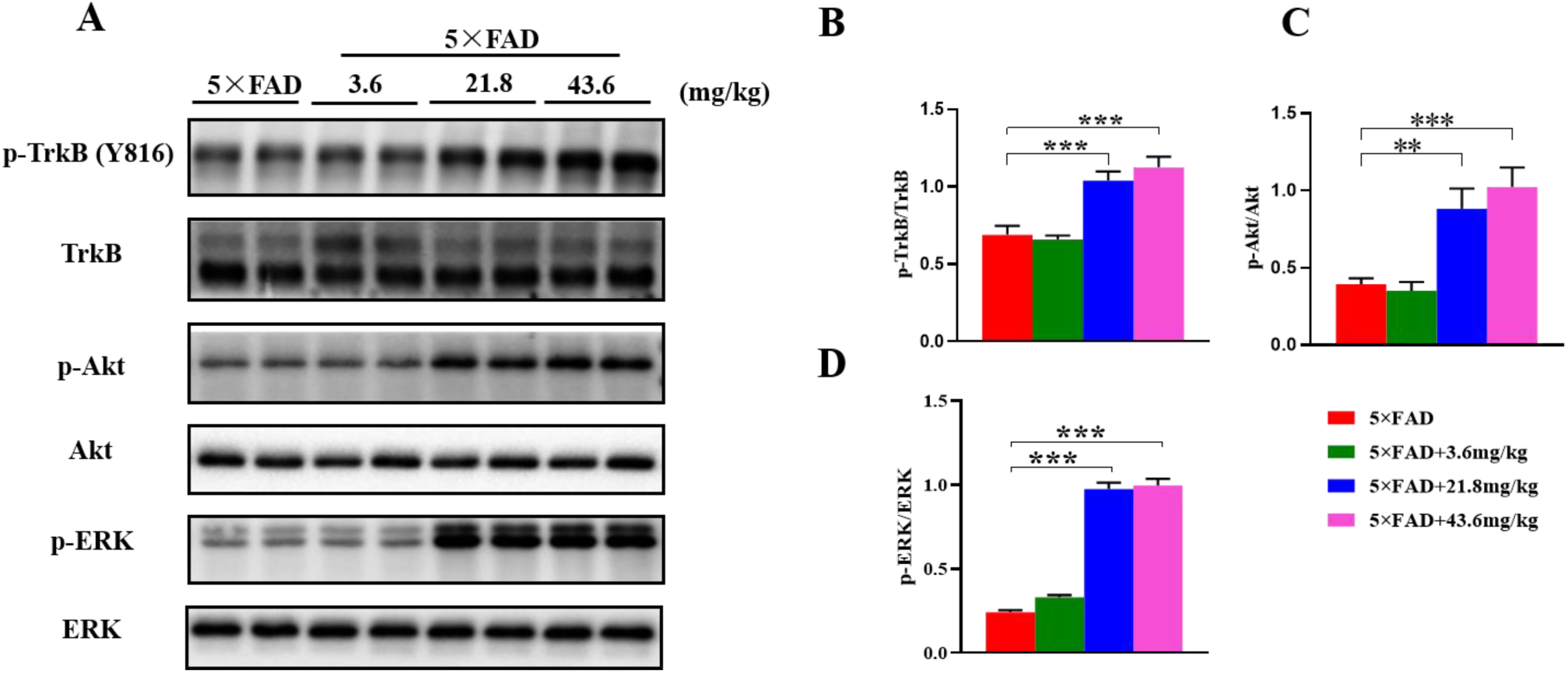
R13 treatment activated TrkB signaling pathway. (A-D) The phosphorylated TrkB at Y816 (p-TrkB) and its downstream signals (Akt, ERK and phosphorylated level) were detected by western blotting (A) and quantitative analyses (B-D). **, *p* < 0.01, ***, *p* < 0.001. All data were expressed as mean ± S.E.M (n=4 mice /per group).

### R13 decreases Aβ levels and tau phosphorylation in 5×FAD mice

AD is mainly characterized by Aβ amyloid deposits and tau-hyperphosphorylation in the brain (Jiao, Yao et al. 2015). Thus, we further detected the deposition of Aβ by immunohistochemistry and dot blots. 5×FAD mice showed evident amyloid deposit in the cortex compared with non-transgenic mice. Interestingly, Aβ amyloid deposit was markedly decreased by R13 in the dose of either 21.8 or 43.6 mg/kg, although low dose 3.6 mg/kg R13 had no obvious effect (Fig. 7A-C). Finally, we detected the effects of R13 on tau phosphorylation in 5×FAD mice. The levels of phosphorylated tau at Ser404 and Ser396 were significantly decreased by R13 treatment, but low dose of R13 revealed no obvious influence in phosphorylated tau in 5×FAD mice (Fig. 7D, E). Notably, no significant changes of total tau expression were observed (Fig. 7D, E).

**Fig 7.**
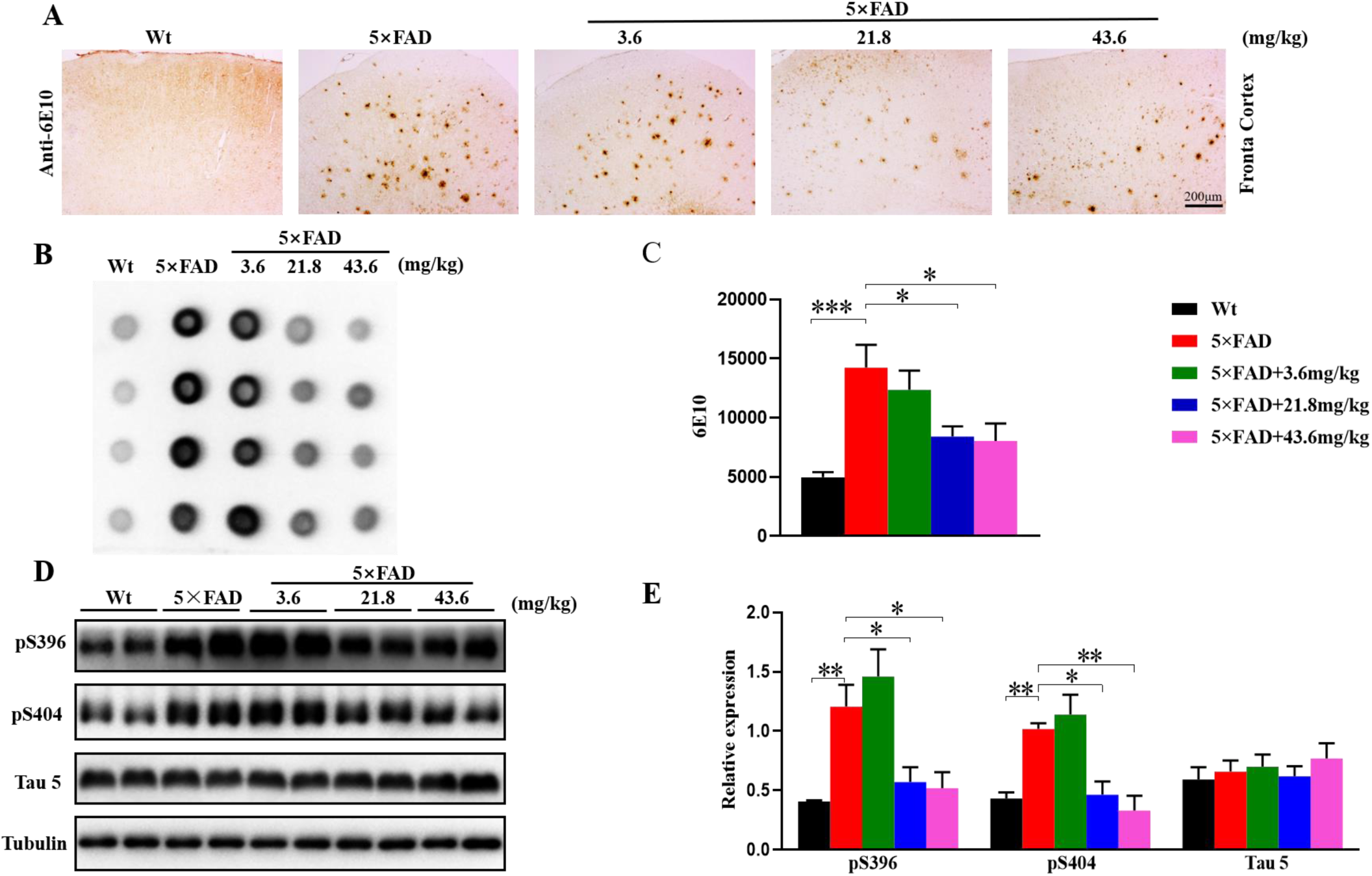
R13 decreased Aβ level and tau phosphorylation. (A) Immunohistochemistry staining for Aβ plaque in frontal cortex of 5×FAD model (scale bar: 200 μm). (B-C) Aβ level in frontal cortex of 5×FAD model was detected by dot blots (B) and quantitative analyses (C) (n=4 mice/per group). (D-E) The levels of phosphorylated tau at Ser396 and Ser404, and total tau (Tau 5) in the hippocampus of 5×FAD model by western blotting (D) and quantitative analyses (E) (n=4 mice/per group). *, *p* < 0.05, **, *p* < 0.01, ***, *p* < 0.001. All data were expressed as mean ± S.E.M.

## Discussion

BDNF-TrkB signaling pathway has been reported to play a key role in learning and memory (Boulle, Kenis et al. 2012). The expression level of BDNF is significantly decreased in the hippocampus of 3-m old 5×FAD mice (Ano, Ohya et al. 2020). R13 is a prodrug of 7,8-DHF, which stably exists in an acidic environment. When R13 reaches the intestine through oral administration, it easily hydrolyzes into an intermediate T1, and finally releases 7,8-DHF in *vivo* (Chen, Wang et al. 2018). Here, we treated 2-m old 5×FAD mice with R13 for three consecutive months. Our results demonstrated that R13 successfully mimicked the effects of BDNF, which activated the TrkB receptor and its downstream signaling pathways including ERK and Akt. By proteomics and western blotting, we found that R13 increased ATP production via activating the OXPHOS signaling pathway and promoted mitochondrial biogenesis. Therefore, R13 significantly improved the spatial learning and memory ability of 5×FAD mice with decreased amyloid plaques and tau hyperphosphorylation.

Mitochondria play pivotal roles in brain functions including energy production, synaptic transmission and cognition (Picard and McEwen 2014). Mitochondrial dysfunction is closely related to the onset and progression of AD (Yang, Li et al. 2017), and energy metabolism disorders are obvious abnormalities in AD patients (Wang, Wang et al. 2014, Cunnane, Trushina et al. 2020). Moreover, compared with wild type mice, 5×FAD model has a significant decrease in ATP levels in the hippocampus (Zaroff, Leone et al. 2015). In the present study, we analyzed the differential proteins located in the mitochondria, and mapped the mitochondrial protein profile by mitochondrial proteomics. As described in the heatmap, complex I, complex III and Complex IV were decreased in 5×FAD model relative to Wt mice, while complex V was increased. However, R13 restored levels of all these proteins in the hippocampus of 5×FAD mice. Then, we verified the proteomics results by western blotting. Complex I (NDUFA10), complex II (SDHB), complex III (UQCRFS1), and complex V (ATP5A) levels all increased induced by R13. These results are basically consistent with the proteomics data. In addition, R13 treatment significantly reversed the decreased ATP levels in the hippocampus of 5×FAD model. These data suggested that R13 corrected energy metabolism disorders in 5×FAD mice. BDNF improved synaptic plasticity by promoting mitochondrial energy production, because BDNF can enhance glucose utilization in the primary neurons in response to energy deficiency (Burkhalter, Fiumelli et al. 2003). Moreover, environmental factors including exercise and dietary calorie restriction promoted BDNF expression to increase energy supply to meet the cellular energy demand (Markham and Greenough 2004). Here, we found that R13 mimicked the functions of BDNF to increase ATP levels by enhancing oxidative phosphorylation.

It has been reported that mtDNA defects have also been related to AD (Coskun, Beal et al. 2004, Markham and Greenough 2004). Moreover, mitochondrial biogenesis is primarily regulated by PGC-1α (Knutti and Kralli 2001, Puigserver and Spiegelman 2003). PGC-1α is highly expressed at neurons because they demand a mass of energy. PGC-1α co-activates and interacts with many transcription factors such as NRF1 (Puigserver and Spiegelman 2003). NRF1 promotes the expression of TFAM, which plays a key role in mtDNA transcription and replication (Lehman, Barger et al. 2000). Furthermore, NRF1 can regulate mitochondrial respiratory complex subunits by controlling the transcription of nuclear genes (Cheng, Hou et al. 2010). BDNF/TrkB signaling activates CREB and AMPK, which are important regulators for PGC-1α expression (Finkbeiner, Tavazoie et al. 1997, Chan, Tse et al. 2015). Our study found that R13 treatment activated AMPK and CREB (pAMPK and pCREB increased), and increased the levels of PGC-1α, NRF1 and TFAM, which suggest that R13 provokes mitochondrial biogenesis in 5×FAD mice via AMPK/CREB/PGC-1α/NRF1/TFAM cascade.

Aβ and tau hyperphosphorylation are two important pathological features of AD. Previous study reported that BDNF reduced tau hyperphosphorylation by TrkB/PI3K signaling pathway (Elliott, Atlas et al. 2005). Moreover, BDNF-TrkB signaling also played an important role in downregulating β-secretase enzyme (BACE1) levels (Devi and Ohno 2012). Here, we show that R13 treatment significantly reduced the levels of Aβ plaque and tau hyperphosphorylation via TrkB activation.

In summary, our present study indicates that R13 activates TrkB to promote mitochondrial biogenesis by activating the PGC-1α/NRF1/TFAM signaling pathway, and improves ATP levels by enhancing OXPHOS. R13 treatment also reduces tau phosphorylation levels and Aβ deposition. All of these contributes to reverse 5×FAD cognitive impairment by R13 treatment (Fig. 8).

**Fig 8.**
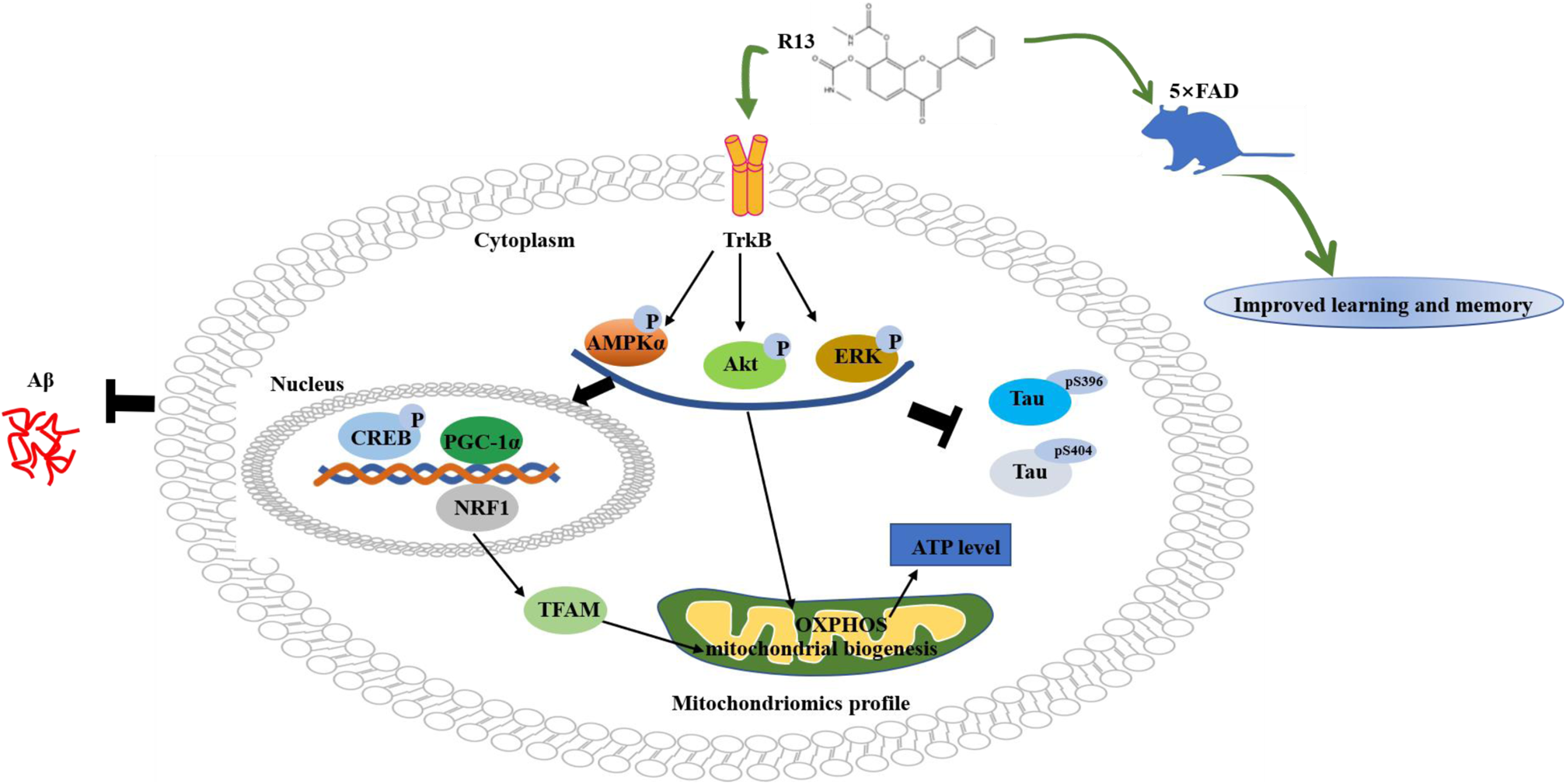
Proposed potential mechanisms by R13, a small-molecule TrkB receptor agonist, improved learning and memory defects of 5×FAD mice. R13 activated TrkB and downstream signaling pathways to promote mitochondrial biogenesis by activating the PGC-1α/NRF1/TFAM signaling pathway, and improved ATP levels by enhancing OXPHOS. Tau phosphorylation levels and Aβ deposition were reduced by R13 treatment. Overall, the diagram illustrated a mechanism by which R13 restored 5×FAD cognitive impairment.

## Data Availability

The data sets used for the current study are available from the corresponding author upon reasonable request.

## Acknowledgements

This work is supported in parts by Natural Science Foundation of China (81673134, 81870846), the Ministry of Science and Technology of China (2016YFC13058001), Guangdong Provincial Key S&T Program (2018B030336001) and Sanming Project of Medicine in Shenzhen (SZSM201611090).

## Author contributions

G.P.L. and X.F.Y. conceived the project, designed the experiments, and wrote the manuscript. X.L. and T.L. designed and performed most of the experiments. H.Y. performed the informatics analysis. S.P., L. Z.J.Z.Y.M.X. and X.R.S. designed, the experiments, analyzed the data and revised the manuscript. J.J.L. performed the proteomics analysis.

## ADDITIONAL INFORMATION

The online version of this article contains supplementary material, which is available to authorized users.

## Supplementary Figure legend

**Supplementary Figure. 1.**
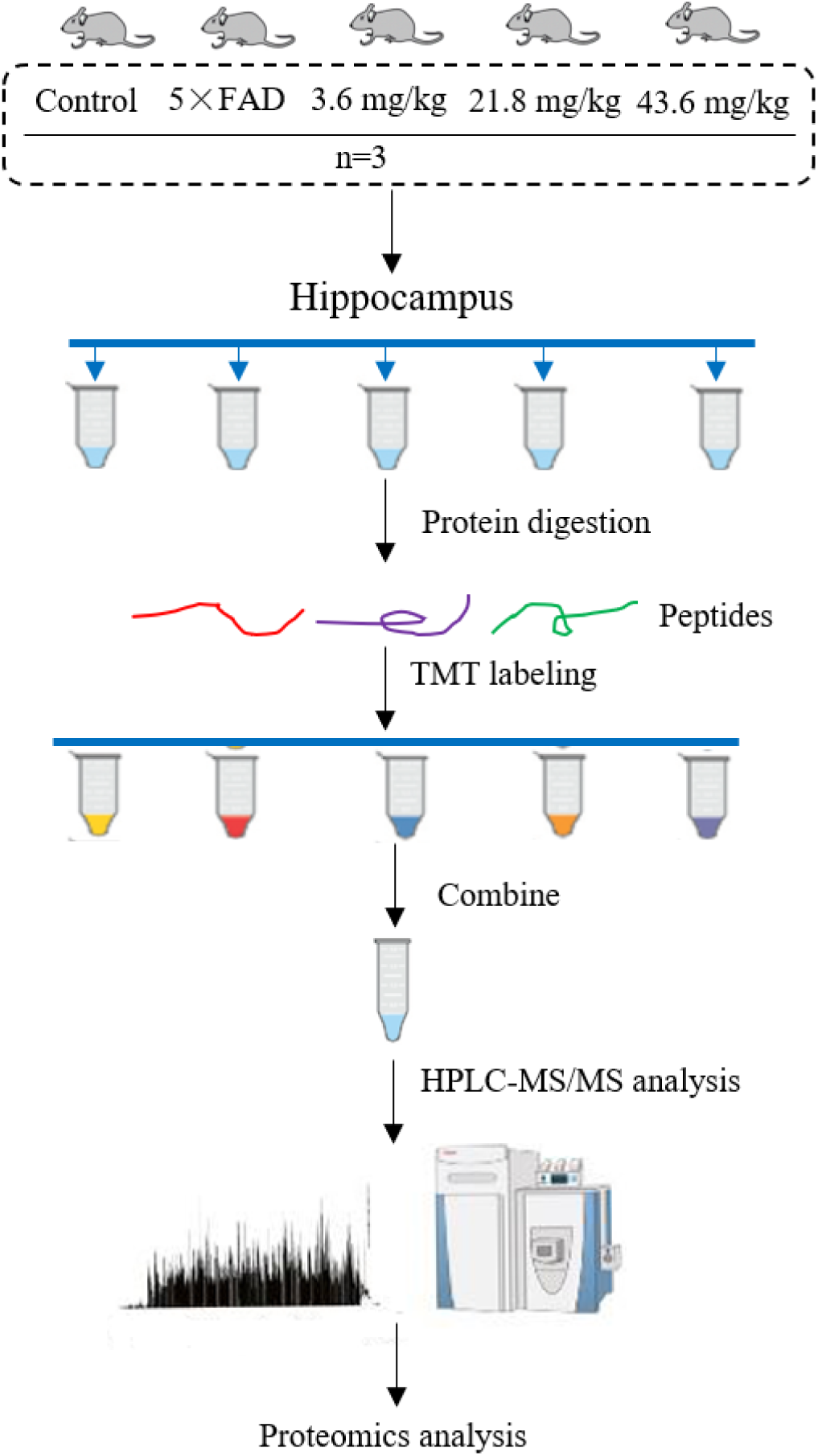
**Schematic diagram of proteomics workflow.**

